# Nanotube mediated cell-to-cell communication and cannibalism in *Halobacillus* sp. GSS1 isolated from Sundarbans, India: A cryptic story of survival under nutrient-limiting condition

**DOI:** 10.1101/2020.10.20.340307

**Authors:** Manas Kumar Guria, Sohan Sengupta, Maitree Bhattacharyya, Parimal Karmakar

## Abstract

Microorganisms play a self-protective role by evolving their genetic and metabolic machinery to thrive in extreme environmental habitats. Halophiles are such salt-loving extremophilic microorganisms able to adapt, survive, and tend to grow at high salt concentrations. In this study, we have isolated *Halobacillus* sp. GSS1 from Sundarbans mangrove, India having a strong salt-tolerant ability (up to 4M) in Zobell Marine 2216 medium. The salt adaptation mechanism of *Halobacillus* sp. was investigated by Confocal microscopy using [Na^+^] specific dye, ‘Sodium Green’ indicating the ‘salt-in’ strategy for their osmoadaptation. Electron microscopic studies revealed that a contact-dependent cell-to-cell communication was profound among the *Halobacillus* sp. under nutrient limiting condition. This communication is mediated by ‘nanotube’, which is highly recommended for the exchange of molecular information between the two individual bacteria. The existence of the ‘ymdB’ gene strongly supports our claim for nanotube formation by *Halobacillus* sp. GSS1. Surprisingly, *Halobacillus* sp. not only utilizing the nanotubes for communication, rather they desperately use nanotubes as a survival weapon under nutrient limiting conditions by triggering cannibalism. This is the first-ever report on the existence of nanotube mediated cell-to-cell communication and cannibalism in any halophilic bacteria, isolated from Sundarbans mangrove forest, India.

**Highlights:**

1. The existence of nanotube mediated cell-to-cell communication was discovered in *Halobacillus* sp. GSS1, isolated from Sundarbans mangrove, India.
2. The communication of *Halobacillus* sp. GSS1 was established through single or multiple nanotubes with the neighboring cells.
3. Intercellular nanotube communication was possible only after the participation of two individual bacteria.
4. *Halobacillus* sp. GSS1 also uses these nanotubes as a survival weapon by triggering the cannibalism to kill their genetically identical siblings.
5. The presence of the ymdB gene in *Halobacillus* sp. GSS1 strongly confers the evidence of nanotube formation.

**Graphical Abstract:** 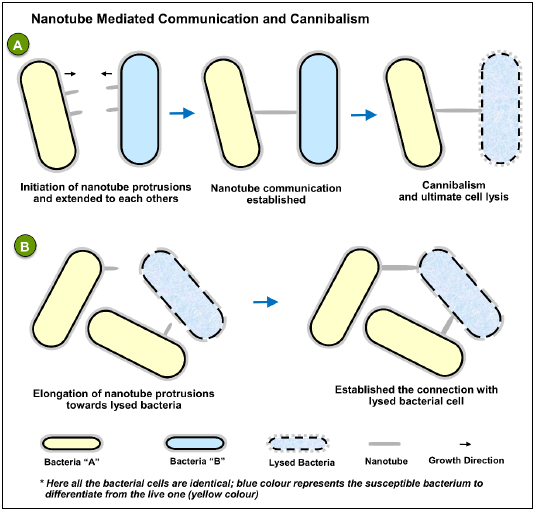

## Introduction

In our living planet, microorganisms are ubiquitously present in the natural environment to extreme microbial habitats. To survive in harsh environments, microbes have evolved their genetic and metabolic activities enabling them to reproduce (Arora and Panosyan, 2019). In this ecological niche, microorganisms are present in a community that helps them to establish an interspecies communication and sometimes they exhibit competitive behaviours for limited environmental space and resources. They can also coexist with other groups of microorganisms to exchange metabolites through various types of physical interaction. These types of interactions among the microbial communities may positively or negatively affect the overall fitness and performance of the individual cells (Benomar et al., 2015, Pande et al., 2015). The interactions between the bacterial species may be through diffusible molecules or direct cell-to-cell communication (Konovalova and Søgaard-Andersen, 2011).

Generally, bacterial communication and the exchange of cellular information occurs through secreted signalling molecules, known as quorum sensing (Long et al., 2009). These molecules can be transferred to the extracellular milieu by membrane diffusion or directly into the cytoplasm of neighboring cells. The secretion of small diffusible signalling molecules allows the bacterial cell to determine the cell density and coordinate their activities (Bassler et al., 2006; Ng and Bassler, 2009; Henke and Bassler, 2004). Sometimes bacteria use these diffusible factors for the intercellular competition to kill their non-resistant and non-immune siblings by secreting microcins, bacteriocins (Destoumieux-Garzon et al., 2002; Cascales et al., 2007; García-Bayona et al., 2019), and toxins (Ellermeier et al., 2006).

Bacteria can also communicate through direct cell-to-cell contact between cells. For instance, conjugation is the first observed classical example of physical interaction in bacteria, where hereditary genetic information is transferred through a type IV secretion system (Lederberg and Tatum, 1946; Alvarez-Martinez and Christie, 2009; Konovalova and Søgaard-Andersen, 2011). Contact-dependent communication in bacteria can also be mediated by various extracellular apparatuses such as Type III, V, and VI secretion systems to deliver the DNA and/or protein effectors that restrain the behaviour of the target organism (Hayes et al., 2010). Moreover, a new mode of cell-to-cell communication was found in bacteria called “nanotube”, which provide an effective conduit to exchange the cytoplasmic constituents between the connected bacterial cells. This interaction may help to provide a synergistic role within and between the microbial communities (Dubey and Ben-Yehuda, 2011; Pande et al., 2015). In contrast, contact-dependent interaction has an antagonistic effect that might be detrimental to other microorganisms by predation or cannibalism, parasitism, and/or sometimes through inhibitory factors or toxin released by cells (Aoki et al., 2005; Hood et al., 2010, Little et al., 2008). This inhibition or killing of neighbouring cells would provide the perpetrator a favourable growth advantage to sustain their life for a longer period.

Halophiles are salt-loving extremophilic microorganisms that can adapt and thrive in hypersaline environments with the requirement of 1 to 5M of NaCl for their growth. *Halobacillus* is a physiologically heterogeneous group of moderately halophilic bacteria, found in saline and hypersaline environments. To maintain the cell physiology and osmotic balance in high salinity, halophiles are evolved two fundamentally different strategies for their osmoadaptation. Firstly, the “salt-in” strategy, where they sequester cations (sodium or potassium) into the cytosol to maintain the ionic gradient across the cell (Thombre et al, 2016; Vaidya et al., 2018). On the other hand, few other halophilic microorganisms use the “salt-out” strategy by excluding salt ions from the cytoplasm, simultaneously synthesizing (*de novo*) and/or accumulating compatible solutes such as amino acids and their derivatives (glycine-betaine, ectoine, proline, glutamine) and the sugars and polyols (trehalose, glycerol, sucrose, sorbitol, mannitol) to maintain the internal osmotic balance. These strategies are largely used in three domains of life viz. bacteria, archaea, and eukarya (Litchfield, 1998; Roberts, 2005; DasSarma and DasSarma, 2015; Thombre et al, 2016).

Sundarbans mangrove is such a type of saline environment, where a variety of halophilic microorganisms belongs to this ecological niche. Sundarbans is the world’s single largest chunks of mangrove forest and only tiger mangrove land situated in the delta of river Ganga and Brahmaputra (Manna et al., 2010). This mangrove ecosystem is unique for its ecological dynamics and rich source in various species diversity including microorganisms (Ghosh et al., 2010). Due to higher salinity, it is suitable to harbour a variety of halophilic microorganisms like halobacteria, haloarchaea, and microbial eukaryotes. Moreover, high index species diversity and limiting nutrient resources compel the indigenous microbial populations to establish an interspecies competition among them. This competition and strong selection pressure have resulted in the evolution of diverse survival strategies within the microbial communities (Sengupta et al., 2015, Sengupta et al., 2020).

In this study, we have isolated a halotolerant bacterium, *Halobacillus* sp. GSS1 from the Bonnie camp, Sundarbans, India. Prior studies on microbial diversity of Sundarbans have been reported (Basak et al., 2015a; Chakraborty et al., 2015; Dhal et al., 2020), but there is no description or clarification on the survival strategy of halophilic bacteria under nutrient challenge. For the first time, we had a remarkable observation in *Halobacillus* sp. GSS1, where nanotube mediated intercellular communication among the bacterial cells were revealed under nutrient limiting condition. Despite their cell-to-cell communication, *Halobacillus* sp. aptly uses these nanotubes as a survival weapon by killing their siblings and feeding on the dead cells. Electron microscopic study verified how bacterial cells initiate the membrane bulging to form a complete nanotube for the cellular bridging. To our knowledge, contact-dependent communication based bacterial survival under saline environment have not been previously reported. Our discovery may provide a new horizon on the adaptation strategy of halophilic bacteria, developed under the extreme environmental condition to survive their lives.

## Results

### Identification and characterization of the halophilic isolate

A total of 4 strains were isolated from Bonnie camp, Sundarbans, among which one strain was selected for further studies based on the salt tolerance ability. The possible growth medium screened for the isolate was Zobell Marine (ZM) 2216 due to the presence of minerals and salt contents that mimics the composition of seawater. The optimum pH for the growth of the isolated bacterium was pH-7.6 (data not shown). Based on the salinity survival in a wide range of NaCl from 1M to 4M, the isolate was screened for further molecular identification. The isolated strain was primarily characterized based on the physiology and morphology as gram-positive (+Ve), rod-shaped, aerobic bacteria (Table 1). The 16s rRNA gene sequencing analysis revealed that the isolated strain SH1 belongs to the *Halobacillus* genus and it exhibits a close evolutionary relationship with *Halobacillus alkaliphilus* FP5 (T) (Fig. 1).

**Fig. 1.**
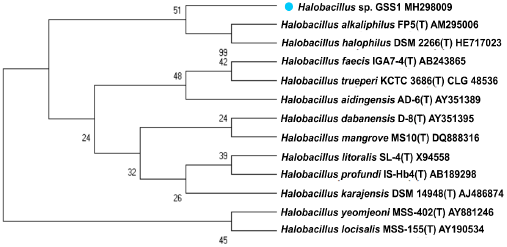
Phylogenetic tree based on 16S rRNA gene sequences obtained by the Neighbor-Joining (NJ) method showing the phylogenetic relationships of the *Halobacillus* sp. GSS1 with the related species. The bootstrap consensus tree inferred from 1000 replicates is taken to represent the evolutionary history of the taxa analyzed. The percentage of replicate trees in which the associated taxa clustered together in the bootstrap test (1000 replicates) are shown next to the branches. The evolutionary distances were. computed using the Jukes-Cantor method and are in the units of the number of base substitutions per site. The analysis involved 13-nucleotide sequences. Evolutionary analyses were conducted in MEGA6. NCBI Accession numbers are given after the species or strain name.

**Table 1.** Morphological and physiological characteristics of the isolated *Halobacillus* sp. strain from Sundarbans, India.

| Isolate | Colony colour | Gram Character | Size of the Bacteria |  | Morphology | Arrangement |
| --- | --- | --- | --- | --- | --- | --- |
|  |  |  | Length (µm) | Width (µm) |  |  |
| SH1* | Yellowish | Gram-positive | 2-3.5 | 0.5-1 | Rod-shaped | Singly or in chains |

\*SH1-Name of the isolate

### Effect of sodium chloride on the growth of Halobacillus sp

The effect of different NaCl concentrations (e.g. 1M, 2M, 3M & 4M) on the growth of *Halobacillus* sp. were performed. Experimental data indicate that the growth profile of isolated strain in the ZM 2216 medium was varied depending on the concentrations of sodium chloride (Fig. 2). The isolate exhibited different growth times of 26 h, 26h, 50h, and 74 h for 1M, 2M, 3M, and 4M NaCl to attain the stationary phase respectively. However, the duration of the ‘lag phase’ increased in 4M in comparison with the other concentrations of NaCl and this may be due to the slower adaptation of bacteria at higher concentrations. The isolated halophilic bacteria showed the range of growth at 1M, 2M, 3M, and 4M NaCl concentration with a generation time of 10.8 min, 15.4 min, 40.4 min, and 15.6 min respectively. These results depict that the *Halobacillus* sp. GSS1 isolated from Sundarbans mangrove can grow over a wide range of NaCl concentrations in ZM 2216 medium.

**Fig. 2.**
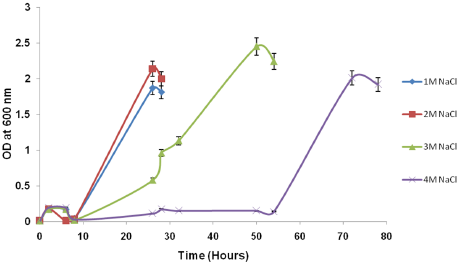
Effect of different concentrations of sodium chloride on the growth of *Halobacillus* sp. GSS1. (Bar represents the standard errors).

### Intercellular [Na^+^] accumulation study using CLSM

To explore the osmoregulation strategy of *Halobacillus* sp. GSS1 isolated from Sundarbans, we have performed a confocal microscopy-based ion imaging detection of intercellular [Na^+^] accumulation study. For measuring the Sodium Green fluorescence intensity inside the cells, the CLSM image was analysed using ImageJ software. Intercellular [Na^+^] ion imaging of *Halobacillus* sp. grown in different NaCl containing ZM 2216 medium revealed that the bacterial cells sequester [Na^+^] ions into the cytosol to maintain the osmotic stress with the external environment. The fluorescence intensity slightly differs with variation in sodium chloride concentrations, notably exception in 4M where the intensity was lower than other concentrations (Fig. 3). This may be due to the leakage of the cell membrane at 4M NaCl.

**Fig. 3.**
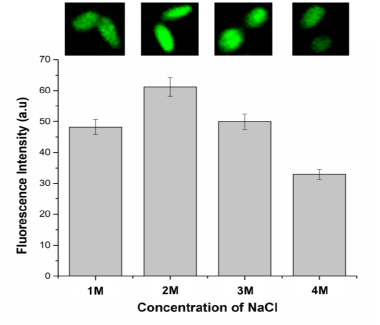
Relative fluorescence intensity of intracellular [Na^+^] ions in *Halobacillus* sp. GSS1 in response to increasing salinity (1M to 4M NaCl). The inset shows confocal microscopy images of bacterial cells incubated at different NaCl concentrations, where [Na^+^] ions fluorescence inside the cytoplasm. (Bar represents the standard errors).

### Electron microscopy for intercellular communication and cannibalism study

The bacterial morphology was investigated by high-resolution electron microscopic studies (SEM & TEM) at the stationary phase of growth under optimal conditions. Scanning electron microscopic images revealed that in each concentration of NaCl (1M to 4M) the bacterial cells exist in a community and few of them were lysed. Interestingly, a small extracellular protrusion appears from most of the cells in every concentration of NaCl containing medium. Two types of the bacterial population were observed from these images, where a small group of the cells does not have the protrusions, but the majority of the cells contain the protrusion on their surface (Fig. 4). Higher magnification of SEM micrographs revealed a cellular bridging between the bacterial cells through tubular protrusions (nanotube) (Fig. 5A & 5B). We observed that a single bacterium concomitantly communicating with more than one neighboring cells through multiple nanotubes (Fig. 5C, & 5D). Besides, scanning electron microscopic study evidenced that, multiple nanotubes are projected from the cell surface at different positions (Fig. 5D, 5F & 5G). Surprisingly, a similar projection of nanotubes in opposite direction was found in the neighboring cell’s surface; perhaps look for communication with neighbours (Fig. 5D, 5E, 5F & 5G). The length of the nanotubes approx. 200 to 500 nm, whereas the width was approx. 90 to 200 nm. However, the length may vary depending on the distance between connected bacterial cells. Higher magnification image evidenced that, few thick nanotubes are appeared other than thin nanotubes; may be due to the need to exchange the larger cargo molecules (Fig. 5F, 6B & 6C) (Dubey and Ben-Yehuda, 2011).

**Fig. 4.**
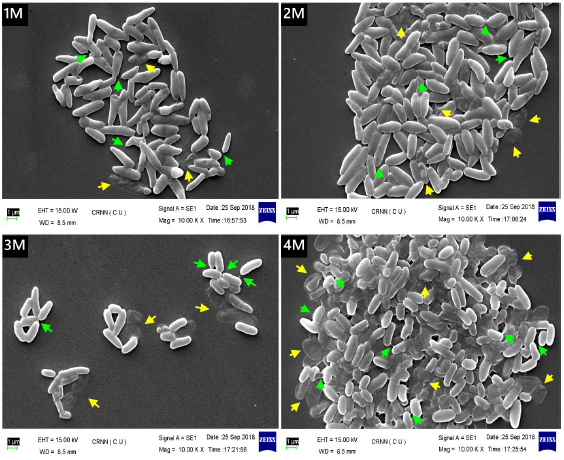
SEM micrograph of *Halobacillus* sp. GSS1 at different concentrations of NaCl (1M to 4M), after harvested from the stationary phase of growth. Bacterial cells tend to communicate with neighboring cells through intercellular nanotube (indicated by green colour arrow). The lysed cells are also observed (indicated by yellow colour arrow). The scale bar represents 1 μm.

**Fig. 5.**
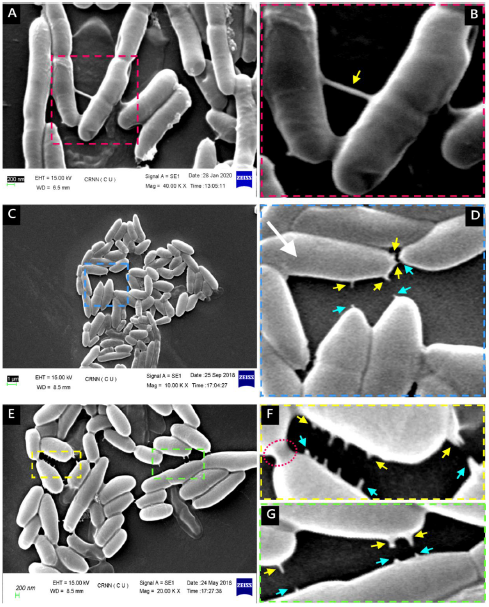
Nanotubes formation and intercellular communication in *Halobacillu*s sp. GSS1. Cells were collected from the stationary phase of growth and observed under SEM. (**A**) A field of *Halobacillu*s sp. community(X 40,000). (**B**) A zoom-in view image of the box region in (A), which revealed a long tubular nanotube connection (indicated by yellow colour arrow) between the bacterial cells. The scale bar represents 200 nm. (**C**) Multiple protrusions emanated by a single bacterium to communicate more than one nearby cells at a time (X 10,000). (**D**) A zoom-in view image of the box region in (C). Two different arrows (yellow and cyan) were used to indicate the extension of nanotubes towards each other for their communication. The scale bar represents 1 μm. (**E**) Multiple projections of nanotubes from both participating cells confirm the involvement in intercellular communication (X 20,000). (**F**) & (**G**) A zoom-in view image of the box region in (E). Two different arrows (yellow and cyan) were used to indicate the extension of nanotubes towards each other for their communication. The red coloured dashed circle indicates the presence of a thick nanotube. The scale bar represents 1 μm.

However, the detailed morphology of nanotube and their projections outlined from the surface of *Halobacillus* sp. GSS1 was also evident by HR-TEM studies. Our analysis revealed that the tubular extensions protruding simultaneously from both neighboring cells and extending towards one another (Fig. 6A to 6F). This result confers the possibility of “bacterial talk” through quorum sensing molecules before protruding the nanotube extension. The initiation of nanotube formation was going through the distortion of the cell membrane to form a “cone-like protrusions” outwardly and continuously extended with time to build a complete nanotube structure (Fig. 6D, 6E & 6F). Notably, the structural arrangement of nanotubes found in *B. subtilis* is composed of “chains of sequential beads” (Dubey and Ben-Yehuda, 2011), which significantly differ from our findings. Multiple distortions on cell membranes were observed to accomplish the subsequent protrusions of nanotube simultaneously (Fig. 6A–6C). The TEM micrographs also revealed the structural resembles of thick nanotubes with SEM, prevailing the communication with their neighboring cells (Fig. 6B & 6C).

**Fig. 6.**
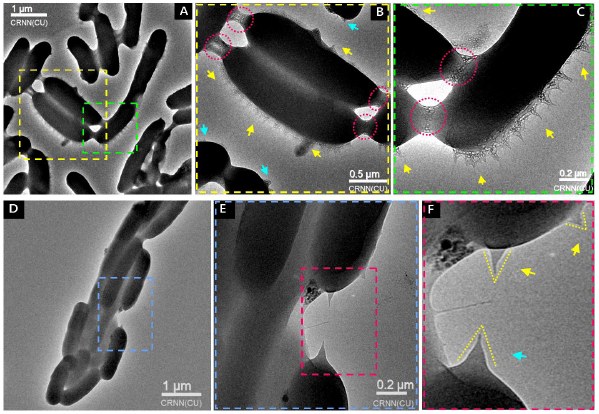
The framework of nanotube formation and cellular bridging in *Halobacillu*s sp. is explicitly observed under HR-TEM. Cells were collected from the stationary phase of growth and processed for TEM micrograph. (**A**) A field of *Halobacillu*s sp. community (Scale bar represents 1 μm). (**B**) A higher magnification image of the dashed square region (yellow) in (A), which revealed the presence of multiple protrusions (yellow arrow) in the cell surface. Cyan arrow indicates the initiation of similar bulging from the neighbouring cells surface. The dashed circle (red) indicates the cellular bridging through thick nanotubes. The scale bar represents 0.5 μm. (**C**) A higher magnification image of the dashed square region (green) in (A), which depicts the multiple “cone-like protrusions” extends outwardly in the cell surface (indicated by yellow arrows). The scale bar represents 0.2 μm. (**D**) The “cone-like protrusions” emanating from the individual bacterial cells for their communication. The scale bar represents 1μm. (**E**) A higher magnification image of the dashed square region (blue) in (D). The scale bar represents 0.2 μm. (**F**) A zoom-in view image of the box square region in (E). Membrane bulging indicates by the yellow and cyan arrows to represents the participation of two nearby cells simultaneously for their communication.

Strikingly, we found an interesting phenomenon using high-resolution electron micrograph studies (SEM & TEM), where *Halobacillus* sp. established a nanotube mediated cellular bridging with neighbouring cells, either live or die. The attachment of nanotube was profound on the core of the cytoplasm of the susceptible cell. No distortion was found on the nanotube and it is well connected in between the neighbouring cells. (Fig. 7A to 7D). However, few healthy cells emanate their nanotubes directing towards the neighbouring lysed cells (Fig. 7E & 7F). Sometimes, these communications were pronounced with multiple cells, may be to overcome the challenging situation like nutrient limitation (Fig. 7G & 7H). As the limitation of the nutrient is profound in the stationary phase, so there may be a nutrient urgency in cells that will force them to engage with more than one partner simultaneously.

**Fig. 7.**
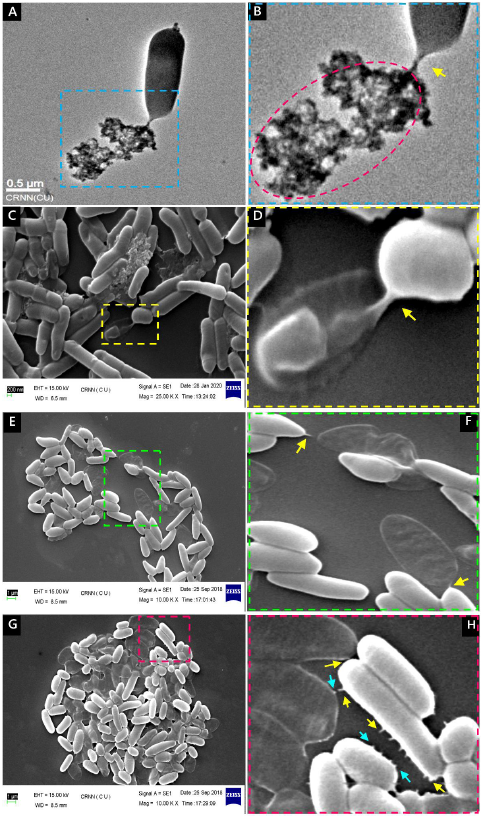
The phenomena of cannibalism in *Halobacillus* sp. GSS1. (**A**) A high-resolution TEM micrograph revealed a connection between the healthy and susceptible cell mediates by nanotube. The scale bar represents 0.5 μm. (**B**) A zoom-in view image of the box square region (blue) in (A) depicts the nanotube link in between the cells (indicated by the yellow arrow). (**C**) An image (X 25000) of nanotube mediated communication with cannibalised bacterium visualized under SEM. The scale bar represents 200 nm. (**D**) A zoom-in view image of the box square region (yellow) in (C) unveiled the attachment of nanotube on the core of the cytoplasm in a susceptible cell. (**E**) SEM micrograph image (X 10000) revealed the nanotube emanating from one cell and try to entangle with nearby lysed cells. (**F**) A zoom-in view image of the box square region (green) in (E). The yellow arrow indicates the protrusions of the nanotube. (**G**) The intercellular connection of bacterial cells with more than one partner including lysed cells occurring simultaneously. (**H**) A zoom-in view image of the box square region (red) in (G). The yellow and cyan arrows indicate the involvement of both the cells for communication.

### Existence of the ymdB gene in *Halobacillus* sp. GSS1

To understand the evolutionary relationship of ymdB gene sequences present in *Halobacillus* sp. GSS1 with the other phylogenetically relevant species, a Neighbor-Joining phylogeny was performed. The phylogenetic analysis revealed that the closest species having ymdB is *Halobacillus litoralis* with 97.32% sequence similarity followed by *Halobacillus faecis*, *Halobacillus trueperi*, and *Halobacillus mangrovi*, sequence similarity ranges between 90-95%. (Fig. 8). The maximum ymdB sequence similarity of nanotube forming *Bacillus* sp. with our strain is only 73%. This evolutionary divergence may bring novel functionality to this metallo-phosphodiesterase, which ultimately contributes to the overall fitness of the organism in a challenging mangrove ecosystem. All of these species have primarily been reported from mangrove ecosystems (Soto-Ramírez et al., 2008) and showed phylogenetic relatedness with *Halobacillus* sp. GSS1 (Fig. 1).

**Fig. 8.**
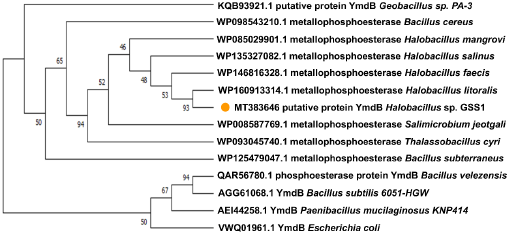
Phylogenetic tree based on ymdB gene sequences showing the phylogenetic relationships of *Halobacillus* sp. GSS1 with their closest species having YmdB. The bootstrap consensus tree inferred from 1000 replicates is taken to represent the evolutionary history of the taxa analyzed. The percentage of replicate trees in which the associated taxa clustered together in the bootstrap test (1000 replicates) are shown next to the branches. This analysis involved 14 amino acid sequences. Evolutionary analyses were conducted in MEGA 6. NCBI Accession numbers were inserted before the name of the strain.

## Discussion

Microbes in the natural environment prefer to live in a community that involves cooperation as well as competition. Microbial interaction and interspecies communication in challenging environments is a relatively unexplored area of research, which has the potential to unveil the true capacity of microbes. A unique environment like the one in Sundarbans, an interface of the marine-estuarine ecosystem is the ideal location to explore the microbial behaviours. Due to the limited consumable nutrient resources and high salinity gradient, a stiff competition within the microbial community can be observed (Basak et al., 2015a; Sengupta et al., 2015). Microbial diversity and their spatial distribution in Sundarbans mangrove have been previously explored (Basak et al., 2015a; Chakraborty et al., 2015; Dhal et al., 2020). Major studies have been reported on the biotechnological potential of the microorganisms isolated from this mangrove ecosystem (Nag et al., 2018; Bhattacharyya et al., 2019). Nevertheless, there is no report on the physiology and the interaction of microorganisms in response to low nutrient resources.

In our study, we attempt to isolate genetically equipped strain, which can withstand the high salinity environment and can manoeuvre their growth under nutrient-limiting conditions. The genus *Halobacillus* belongs to the family Halobacteriaceae and was first described by Spring *et al.* (1996). They are a physiologically heterogeneous group of microorganisms present in saline and hypersaline environments. Our isolate, *Halobacillus* sp. GSS1 exhibited the growth advantage in a wide range of salinity (1M to 4M NaCl) and their growth pattern significantly altered with the change of salt concentrations in the ZM 2216 medium. The stationary phase attained by *Halobacillus* sp. GSS1 was interestingly varied depending on the NaCl concentrations of medium (e.g. 26 h, 26h, 50h, and 74 h in 1M, 2M, 3M, and 4M NaCl respectively). The influence of NaCl concentrations allows the *Halobacillus* sp. to grow at different rates, signifies the physiological adaptability and endurance of the isolated strain in saline environments. We have targeted the stationary phase for all the morphological investigations because of its close mimic to the Sundarbans in-situ condition where a constant competition took place across the microbial community due to the high species richness and limited availability of nutrient. Thus, from this study, it can be assumed that the strain *Halobacillus* sp. GSS1 acquired certain physiological and morphological adaptation to withstand the nutrient-deficient condition at the stationary phase.

To compensate for the high osmotic pressure, halophiles are generally followed two osmoregulation strategies. The classical adaptation strategy e.g. ‘salt-in’ strategy, by sequestering cations (sodium or potassium) into the cells and secondly the accumulation of ‘compatible solutes’ for maintaining the osmotic balance to provide the cellular integrity. It has been reported that *Halobacillus* sp. accumulates a cocktail of different compatible solutes (Hänelt et al., 2013; Saum et al., 2008) as well as adopt a salt-in strategy (Vaidya et al., 2018) to maintain the cellular osmotic balance under salinity stress. However, the *Halobacillus* sp. GSS1 from Sundarbans prefers the salt-in strategy; suggest that cellular osmoadaptation may depend on the ecological environment, where they reside.

In harsh environmental conditions, possible manipulation of cell physiology and behaviours perceives in bacteria to cope with the cellular perturbations. Bacteria have evolved various survival strategies including biofilm formation, secondary metabolite production, bioluminescence, conjugation, and sporulation for their adaptation, which are often controlled by quorum sensing molecules (Wong et al., 2020). For instance, in response to nutrient starvation, marine bacteria and few haloarchaea persist for a long time by altering their behaviours or cellular morphology (Hood et al., 1986; Norton et al., 1988). Besides, environmental stress like desiccation insists to adapt the *Salmonella* sp. by altering their morphology and cell-to-cell communication for long time survival (Habimana et al., 2014). Our study focused on the aspects of cell-to-cell communication in the context of an ecological perspective, particularly in nutrient starvation conditions. We have attained a concrete proof of this notion from the electron microscopic studies, where we found the existence of a unique and unusual cell-to-cell communication system in *Halobacillus* sp. GSS1 connecting adjacent cells. This communication is mediated by membrane-derived long tubular protrusions called “nanotube”. Our investigations suggest that intercellular communication by nanotube is resulted through the participation of different cells, not by a single bacterium. This phenomenon firmly differs from the communication strategy of *Bacillus subtilis*, where nanotubes are emanating from a single cell only (Dubey and Ben-Yehuda 2011). The nanotube formation under nutrient starvation may likely be necessary for nutrient exchange. Benomar et al. (2015) evidenced that two phylogenetically distant co-culture cells exchange their cytoplasmic molecules when the cells have a nutrient shortage. Besides, multiple protrusions were observed by a single bacterium depicts the nutrient urgency and the aggressiveness of cells for nutrition. This might be triggering the cannibalism of susceptible bacteria by inducing the killing factors and inevitably reclaim their nutrient requirement from dead siblings to access the growth advantage for long time survival. We have seen many live cells attached to the dead cell through nanotube probably indicating the utilization of dead cell remnants as a source of nutrients. Thus, it is tempting to speculate that initially two or more live cells may be connected during stress; later few of the cells either may be dead by the stress itself or may be killed by the relatively healthy bacteria through the secretion of some inhibitory factors. The live cells may encode an immunity protein, which can shield themselves to prevent auto-inhibition as has been reported in bacteria (Ellermeier et al., 2006; Hayes et al., 2014; Koskiniemi et al. 2013). Such cannibalism was first observed in *Bacillus subtilis*, where they competing with their siblings for nutrient resources during starvation and delay the sporulation by cannibalising its siblings through secreting extracellular killing factors (Gonzalez-Pastor et al., 2003; Ellermeier et al., 2006).

YmdB is a part of the large calcineurin-like phosphatase/phosphodiesterase family present in all domains of microbial life. According to Diethmaier et al., 2014, YmdB plays a crucial role in regulating late adaptive responses of *B. subtilis*, especially in biofilm formation. Although the identification of the primary target that is subject to YmdB-mediated hydrolysis is still to be deciphered. Surprisingly, a recent study discovered that in *B. subtilis*, a conserved YmdB presents in nanotubes and is essential for both nanotube production and intercellular molecular trade (Dubey et al, 2016). Thus to reinforce our claim for nanotube formation by *Halobacillus* sp. GSS1, we made a successful attempt to find the presence of the ymdB gene in this strain. In this context, it is noteworthy to mention that though the presence of YmdB has been found in various *Halobacillus* sp. whole-genome deposits, there is no morphological evidence of nanotube formation was reported. Although none of the reports was concerned about the connection between YmdB protein and nanotube formation in *Halobacillus* sp. From this viewpoint, this is the first-ever report on “Halobacterial nanotube” by any *Halobacillus* sp. isolated from Sundarbans, India. Therefore, the existence of the ymdB gene does emphasize the presence of nanotube in *Halobacillus* sp. GSS1. All the closely related species who have ymdB sequence similarity with *Halobacillus* sp. GSS1 is primarily reported from the mangrove ecosystem (Fig 8). It can be speculated that the presence of YmdB protein in *Halobacillus* species is frequent and particularly in mangrove habitats, therefore the nanotube formation may be a common phenomenon in those organisms. However, the exact role of bacterial nanotube in a mangrove ecosystem is a future prospect of research. The presence of ymdB in *Halobacillus* sp. and consequently nanotube formation may have a correlation, which triggers the cell physiology in nutrient starvation conditions. However, the expression of YmdB in *E. coli* is transcriptionally activated when the bacterial cells enter into the stationary phase (Kim, et al., 2008), corroborating our observation that the stationary phase emphasis the nanotube formation in *Halobacillus* sp. GSS1 under nutrient limitation. This is the first-ever report on the existence of contact-dependent intercellular bacterial communication mediated by nanotubes in any saline environment to combat nutritional stress. Our study claims that bacterial communication in a saline environment does not only depend on the diffusible quorum sensing molecules but a physical mode of communication among the bacterial communities also prevails. This discovery also helps us to explore the communication pattern of *Halobacillus* sp. with other groups of indigenous microorganisms present in saline environments. Besides, inhibition of bacterial growth through cannibalism might be new hope for promising alternative of antibiotic treatment in the future to control the bacterial pathogenesis.

## Materials and Methods

### Sample collection and isolation of Halophilic bacteria

Soil samples were collected from the Bonnie camp area (21°84′ 26.32′′N 88° 60′ 33.34′′E) Sundarbans, India in sterile falcon tubes and preserved the samples in a cooler at 4°C until further use. All the samples were processed in the lab within 24 hrs after sampling. To isolate the halophilic bacteria, soil samples were then dissolved in 0.9 % saline water and serially diluted (10^-1^ to 10^-6^). 100 uL aliquots from each dilution were plated on different concentrations of NaCl containing Zobell Marine Agar 2216 (Gutierrez T et al., 2020) (HiMedia, Mumbai, India) medium and incubated at 37^0^C until the colonies appear. Sub-culturing the isolates for further characterization and stored as glycerol stocks in a cryovial at −80^0^C for long time preservation. (The composition of the Zobell Marine (ZM) 2216 medium are (g/L) Peptone-5.000, Yeast extract-1.000, Ferric citrate-0.100, Sodium chloride-19.450, Magnesium chloride-8.800, Sodium sulphate-3.240, Calcium chloride-1.800, Potassium chloride-0.550, Sodium bicarbonate-0.160, Potassium bromide-0.080, Strontium chloride-0.034, Boric acid-0.022, Sodium silicate-0.004, Ammonium nitrate-0.0016, Disodium phosphate-0.008, Sodium fluorate-0.0024, pH-7.6±0.2).

### Screening of Halophilic bacteria

The best possible growth medium for halophilic bacteria was optimized in three different mediums such as Nutrient Broth (NB), Luria Bertani (LB), and Zobell Marine (ZM) 2216. However, no growth was observed on LB and NB medium except ZM 2216. Further, the growth of halophilic bacteria were optimized in ZM 2216 medium supplemented with different concentrations of NaCl externally, ranging from 1M, 2M, 3M, 4M and 5M (e.g. 5.84%, 11.68%,17.53%, 23.37% & 29.22% (w/v) respectively) to check the maximum NaCl tolerance capacity. Based on the survival in a wide range of salinity up to 4M NaCl, one isolate was screened from four different isolates. The pH of the ZM 2216 medium was varied (pH-5.0, 7.6, 9.0) by using dilute HCl or NaOH to optimize the pH value.

### 16sRNA gene sequencing and phylogenetic analysis of Halophilic bacteria

The molecular identification of isolated bacteria was performed based on the 16S rRNA gene sequencing method. Total genomic DNA was extracted using the High Pure PCR Template Preparation Kit (Roche, Germany) according to the manufacturer’s instructions and the 16S rRNA gene of the isolate was amplified by PCR using 27F:AGAGTT TGATCMTGGCTCAG (Wilmotte et al., 1993) and 1492R:CGGTTACCTTGTTACGACTT (Lane, 1991) primers. The PCR product is around 1500 bps. The PCR amplicon was purified using the High Pure PCR Product Purification Kit (Roche, Germany) to remove the contaminants and the amplicon was sequenced with forward and reverse primers using Big Dye Terminator v3.1 Cycle Sequencing kit on Applied Biosystems 3730 x l Genetic Analyzer. The consensus sequence of the 16S rDNA gene was generated from forward and reverse sequence data using aligner software. The sequence was initially compared using the BLAST algorithm (Altschul S.F et al., 1997) using the NCBI GenBank database to identify the most similar sequence of phylogenetic neighbours. Based on the maximum identity score, the first ten sequences were selected from the EzTaxon server (Chun et al., 2007) and aligned using multiple alignment software program ClustalW. The distance matrix was generated and the phylogenetic tree was constructed by the Neighbor-Joining method, with 1000 bootstrap analysis using MEGA6 (Molecular Evolutionary Genetics Analysis) (Tamura et al., 2013). The 16S rRNA gene sequence of *Halobacillus* sp. GSS1 was submitted to NCBI GenBank with accession number MH298009.

### Effect of NaCl on the growth of *Halobacillus* sp

The effect of sodium chloride on the growth of halophilic bacteria was explored by inoculating an equal amount of culture in each ZMB 2216 medium, containing four different concentrations of NaCl (e.g. 1M, 2M, 3M &4M). After inoculation, culture flasks were kept in a shaking incubator (130 rpm) at 37^0^C until the growth occurs. The growth of the bacterium was measured in different time intervals at 600 nm absorbance using a UV-visible spectrophotometer (Varian Cary 50 Bio).

### Detection of [Na^+^] uptake capacity by *Halobacillus* sp. GSS1

The qualitative analysis of intracellular sodium ions [Na^+^] accumulation was performed using sodium ion specific fluorescence indicator dye, Sodium Green^TM^ (Invitrogen, USA) (Amorino et al., 1995). The bacterial cells were harvested by centrifugation (5000 rpm, 5 min) from four different concentrations of NaCl (e.g. 1 M, 2 M, 3M, & 4M) containing ZM 2216 medium followed by thorough washing and resuspended in a sodium motility buffer (10 mM potassium phosphate, 85mM NaCl, 0.1 mM EDTA, pH-7.0). To increase the permeability of the cell membrane, bacterial cells were suspended in freshly prepared high EDTA motility buffer (sodium motility buffer, 10 mM EDTA) for 10 min. After that, the cells were thorough washed three times in sodium motility buffer and resuspended the cells (10^8^ cells/ml) in Sodium Green loading buffer (sodium motility buffer, 40 μM Sodium Green^TM^). The samples were then incubated at room temperature for 30-45 min in dark with gentle agitation intermittently. After incubation, cells were thorough washed and resuspended in sodium motility buffer to remove the excess Sodium Green, which was noncovalently associated with the membrane. The cells were then mounted on a glass slide and observed under the Confocal Laser Scanning Microscope (Olympus FV 1000, Model IX81, Singapore). Sodium Green dye was excited in epi-fluorescence mode at 488 nm by an Ar-ion laser (Alexa Fluor) for selective detection of Sodium Green^TM^ inside the cells and observed the image under 100X. Later the images acquired by CLSM were analysed using Olympus FV1000 software.

### Scanning Electron Microscopy analysis

To explore the bacterial morphology at the stationary phase, cells were collected from each ZM 2216 medium containing different concentrations of NaCl. After that, cells were centrifuged at 5000 rpm for 5 min followed by thorough washing with PBS (0.1M, pH-7). The cells were then fixed with 3% glutaraldehyde (Merck, Germany) in sodium phosphate buffer (0.1 M, pH 7.2) for 2 h at 25^0^C followed by overnight incubation at 4^0^C. Next, the cells were postfixed by incubation with 1% osmium tetraoxide for 1h at 25^0^C in the dark. Then the samples were thorough washed with sodium phosphate buffer and gradually dehydrated using a graded series of ethanol (30%, 50%, 70%, 80%, 90%, and 100%). The time of incubation in each solution was 10 min and finally, 1hr in 100% ethanol (Guria et al., 2014). Samples were mounted on aluminium stubs and the cells were sputtered with platinum using a sputter coater and observed at 15 kV with a scanning electron microscope (ZEISS EVO-MA 10, Germany).

### Transmission Electron Microscopy analysis

To study the morphology of *Halobacillus* sp. GSS1, cells were collected from the stationary phase and washed thrice with PBS (0.1M, pH-7) by centrifugation (5000 rpm, 5 min). The bacterial cells were then fixed with 3% glutaraldehyde (Merck, Germany) in sodium phosphate buffer (0.1 M, pH 7.2) for 2 h at 25^0^C followed by overnight incubation at 4^0^C. The fixed cells were washed thrice with PBS (0.1M, pH-7) by centrifugation, and subsequently, the samples are drop cast on a 400-mess TEM copper grid. After draining off the excess amount of samples by blotting with filter paper from the edge of the grid, the samples were immediately stained with 2% aqueous uranyl acetate. The excess amount of stain was discarded by blotting with filter paper and the grids were allowed to air dry. Samples were observed under a high-resolution transmission electron microscope (JEOL JEM-2100 HR, Japan) at an accelerating voltage of 100 kV.

### Identification of the presence of the ymdB gene in *Halobacillus* sp. GSS1

To identify the presence of the ymdB gene in Halobacillus sp. GSS1, we have designed the ymdB specific primers ymdBF:ACTGCTTCTCGATCCATACC and ymdBR:GCATCAGGGAAAGGGATTAC using clone manager software. The PCR product to be amplified is around 515 bps. The ymdB gene amplification was carried out for 35 cycles in a thermocycler (Gene AmpR PCR System 9700, Applied Biosystems, USA) using the following conditions: after 5 min of initial denaturation at 95^0^C, each cycles consisting of denaturation at95^0^C for 30 sec, primer annealing at 50^0^C for 30 s, and primer extension at 72^0^C for 45 sec, followed by a 7 min of final extension step at 72^0^C for the last cycle. Negative control was performed without a template to ensure no contamination occurs. The PCR product was purified using the High Pure PCR Product Purification Kit (Roche, Germany) and subjected to electrophoresis in 1% (w/v) agarose gels to confirm the PCR. The purified amplicons were sequenced in both forward and reverse directions using Big Dye Terminator v3.1 Cycle Sequencing kit on Applied Biosystems 3730 x l Genetic Analyzer (Applied Biosystems, USA). The ymdB gene sequences were compared with existing sequences from the NCBI GenBank database using the BLAST algorithm (Altschul S.F et al., 1997) to identify the most similar sequence of phylogenetic neighbours. The evolutionary history was inferred by using the Maximum Likelihood method and the JTT matrix-based model (Jones et al., 1992). The bootstrap consensus tree inferred from 1000 replicates (Felsenstein 1985) is taken to represent the evolutionary history of the taxa analyzed. Initial tree(s) for the heuristic search were obtained automatically by applying Neighbor-Join and BioNJ algorithms to a matrix of pairwise distances estimated using the JTT model, and then selecting the topology with superior log likelihood value. This analysis involved 14 amino acid sequences. There were a total of 159 positions in the final dataset. Evolutionary analyses were conducted in MEGA 6 (Tamura et al., 2013). The ymdB gene sequence of *Halobacillus* sp. GSS1 was submitted to NCBI GenBank with accession number MT383646.

## Acknowledgments

The author (MKG) would like to acknowledge the DST-SERB National Postdoctoral Fellowship (NPDF) scheme and UGC DS Kothari Postdoctoral Scheme, Government of India for their research funding and support. We are grateful to Prof. Sanjay Ghosh (University of Calcutta) and Dr. Abhrajyoti Ghosh (Bose Institute) for their insightful discussions during this study. DBT-IPLS & Department of Biochemistry (University of Calcutta) for the instrumental support during experiments and CRNN, Kolkata for Electron Microscopy imaging facility. We are thankful to Prothyush Sengupta and Urmila Goswami for their technical supports during EM studies.

## Authors Contributions

M.K.G. and P.K. designed the research plan; M.K.G. performed the experiments with supervision from P.K. and M.B.; M.K.G., P.K., and M.B. wrote the manuscript; S.S. helps to design the primers for ymdB gene, identification, and corresponding information. All authors reviewed the manuscript and gave final approval for publication.

